# Fatigue behavior of trabecular bone orientation

**DOI:** 10.1101/2020.02.12.945352

**Authors:** Fatihhi Szali Januddi, M.N Harun, Jaafar Abdullah, Mohammad Mostakhdemin, Ardiyansyah Syahrom

## Abstract

The present study reports the anisotropy effects of uniaxial and multiaxial loading on cancellous bone in order to mimic true physiological conditions as well as pathological reactions and thereby provides improved data that represents clinical and real life conditions. Cancellous bone samples were CT-scanned for morphological analysis and model construction. The models were then computationally loaded on three different directions; horizontal, vertical, and at 45°. Lower BV/TV, Tb.Th, and Conn.D resulted in lower number of cycles to failure, regardless to the loading conditions. However, the number of cycles to failure was found to be negatively correlated to the value of structural model index. Dramatic increased in effective plastic strain and decrease in cycles to failure were demonstrated by the cancellous bone models under multiaxial loading. The reduction of fatigue life was five times lower in multiaxial condition in comparison to the fatigue life under uniaxial loading. Off-axis orientation effect on the fatigue life of the trabecular bone was demonstrated the worst in horizontal trabecular bone model. Effective plastic strain was recorded the highest in horizontal model, while the model at 45° demonstrated 1.6 times higher effective plastic strain than the vertical ones. This is due to several numbers of thin trabeculae which are susceptible to fatigue at higher stress concentration. In conclusion, the anisotropic effect of uniaxial and multiaxial loading onto the mechanical behaviour of bovine cancellous bone was demonstrated throughout this study. It is apparent that multiaxial with off-axis forces are important to be considered as the loading direction manifests the fatigue lifetime of cancellous bone.

## 1. Introduction

For over 30 years biomechanics research has been widely explored with special interest is sending forth on the influence of trabecular bone towards weakening and failure of whole bone, and how the stimulating remodelling process helps in retaining the bone strength. Clear understanding of the biomechanics of bone is well related in diagnosis and treatment of medical issues such as osteoporosis, bone fracture, bone remodelling, and implant system.

Failure in most loaded biological structures has been characterized as fatigue-induced [1]. Fatigue can be defined as the weakening of a bony material resulted from repetitive applied stresses or strains with accumulation of damage [2]–[6]. Fatigue failure in bone has been found to be resulted from worsened deposition or mineralisation of bone matrix, or the unrepaired microdamage accumulation from daily repetitive loads which increase bone fragility [7]. As fatigue failure in bones contribute to significant clinical implications, studies and investigations to better comprehend fatigue failure in bones are required. Factors affecting fatigue strength of bone include the loading mechanism, frequency, strain rate, age, anatomic site, stiffness, density and temperature, as well as the microstructure of the bone [3], [5], [8]–[11].

With advancement of technology, direct quantitative morphological analysis on three-dimensional (3D) reconstructions is made possible with micro-computed tomography (micro-CT). The morphological indices include volume fraction (BV/TV), trabecular thickness (Tb.Th), trabecular separation (Tb.Sp), and trabecular number (Tb.N). As material testing on highly inhomogeneous structures like the trabecular bone is quite complicated and no standard for experimental conditions are available in terms of sample size, loading rate, loading mode, and surrounding media, results from literature are also diversified. Some of the variations in mechanical data may be ascribed to experimental effects, introduced by ignoring the structural anisotropy, the proper boundary conditions (e.g. end artefact errors) [12] and size effects [13]. But there is also a natural heterogeneity which complicates the analysis of trabecular bone and large variation in between samples properties scattered the results in mechanical interpretation especially in bone mechanic study such as creep or fatigue [14].

Progressive collapses of the vertebrae [15] and loosening of implants [16] have been associated to the damage and creep strain which attracts interest in understanding the associated failure. The number of cycles to failure of the trabecular bone is in direct relationship with the volume fraction, fabric, and applied stress [17]. Lifetime of the trabecular has also been recognised to be influenced by loading direction [4]. Current fatigue assessment on trabecular bone is focused more on the effect of uniaxial compression. This method however leaves out other contributing factors to the strength of bone such as the morphological information. While physiological and traumatic loading are multiaxial in nature, uniaxial assessment limits the reliability of the yielded information. Multiaxial loading demonstrates mixed-mode failure where the damage propagated from one mode (tension) to another (shear). In bioengineering, multiaxial criterion provide better understanding on the relationship of the trabecular tissue structure and its physiology in which will improved implant system and development of bone analogue [18]. To the author’s knowledge, none of the reported works in the literature has ever quantified the behaviour of the trabecular under multiaxial fatigue based on its anisotropy. Thus, this study aims to investigate the influence of anisotropic of cancellous bone under effect of multiaxial loads and how severe the off-axis can affect the fatigue life of this type of bone. Therefore, the outcomes of this present work is hope to shed lights on a few aspects involved in the failure of the trabecular under uniaxial as well as multiaxial loads and contribute information for future development.

## 2. Materials and Methods

### Sample Preparation and µCT Imaging

Trabecular bone samples were extracted from the hind-limb of fresh bovine cadavers gathered from a local slaughter house, then cut from the femoral ball using a ±150 rpm diamond saw (Behringer GmbH, typeSLB 230 DG HA, Kirchardt) under copious lubricant irrigation to minimize heat generation and strut breakage. Saline water was used as a lubricant to ensure that the temperature did not exceed 46 °C to protect the sample from heat-related damage [11]. An infrared thermometer (Fluke 62 Mini Infrared Thermometer) was used in order to observe the temperature of the sample based on the blade and coring bit. During the cutting and drilling procedure, the process was stopped at several stages to measure the temperature. The machining process began by thawing the bone to room temperature. Excision lines were marked (Figure 1) on the bone surfaces to aid the sectioning process from medical-lateral condyle region in distal femur, femoral head and greater trochanter in the proximal femur. These lines were corresponded to the axis of the femur. The sectioning process was done by hand-saw and finished using a precision cutter (Techcut 5™ Precision Sectioning Machine, Allied High Tech, USA) at speed ±150 rpm with diamond wafering blade and continuous lubrication. The sectioned bone was then drilled with 1.55 mm thick diamond coring bit into the trabecular cylindrical samples at 150-250 rpm. The angle of the coring bit used was made sure not to exceed 10° from the principle anatomical axis [19].

**Figure 1:**
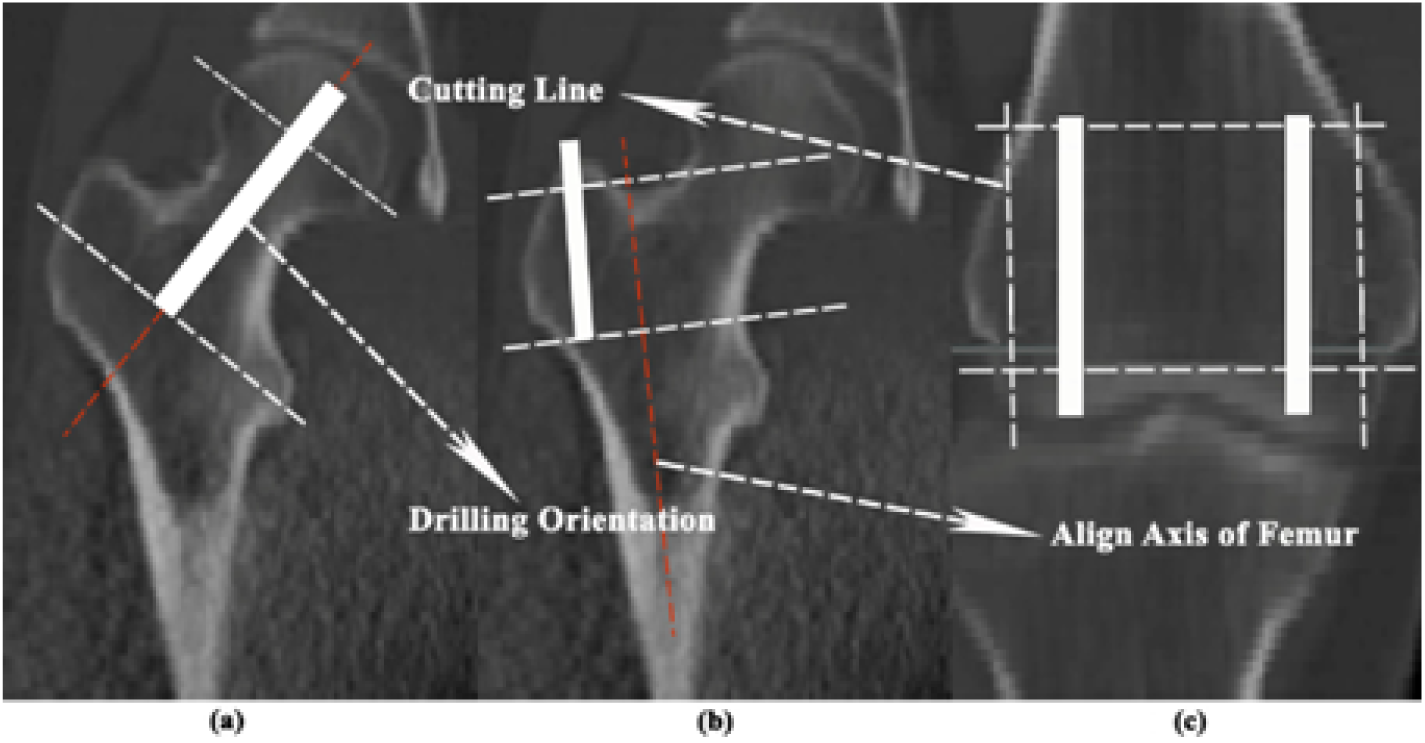
Excision lines on the femur oriented to the bone axis. This figure is a representation only.

Cylindrical samples with a total length of 15 mm and a diameter of 10 mm were then drilled using a 1.5 mm thick diamond-tip coring bit [20]. An electronic calliper was used to measure the length and diameter of the specimens. The cylindrical bone samples were ultrasonically cleaned (Cest ultrasonic, model P1100SR, USA) [20] with a chemical detergent (Pumicized, Gent-l-kleen, USA) [21] for marrow removal. Excessive marrow was further cleansed with water and air jets. The prepared samples were stored in plastic bags and frozed at -20 °C [22]. After at least 24 h, the samples were then prepared and set up for CT scan (Skyscan 1172, Kontich, Belgium). The micrograph of the sample determined its alignment with the principal trabecular direction. A 3D model with a diameter of 5 mm and a length of 5 mm of the trabecular bone was extracted from the central part of the prepared sample and scanned at high resolution of 20 µm. Despite the small size used, the 3D model satisfied the continuum assumption for trabecular bone which is at least to have three to five inter trabecular lengths in size [23]. This is due to the limitation in computational power used in this study that could not possibly construct the whole trabecular model. To study the effect of multiaxial loading based on physiological loading condition, sample of trabecular bone were loaded by multiaxial fatigue and compared to those loaded by uniaxial fatigue of the same sample. Nonlinear simulation was used which included plasticity in stress-strain curve as basis in modelling fatigue under LCF and HCF effects [3]. This modelling is normally developed as strain-based approach [24], [25] and thus can be applied directly to estimate fatigue failure of trabecular bone. The crucial aspect of this approach is to compare the effect of uniaxial and multiaxial loading on the same FE model. The loading values were chosen from normal walking condition taken from hip contact forces where uniaxial loading is defined as the vertically oriented force (z-direction only) and multiaxial loading considers all axes of forces of the gait loading.

### Finite Element (FE) Analysis of Fatigue Loading

The FE model was analysed using the commercial COMSOL Multiphysics FE software (COMSOL Multiphasic software, Burlington, USA, version 3.4). The cylindrical axis was assigned the z-direction for axial fatigue load. Multiaxial loading from three axes contributed to the combined axial and torsional moment. The deformation behaviour of bovine bone has been found to be similar to that of human bone [9]. This study used trabecular bone of the bovine femoral hip as a material representation. In human, this site experiences more multiaxial load transfer through body weight than bone or joints at other sites [26]. The nonlinear-elastoplastic material behaviour of the trabecular bone was modelled with the following values: 1000 MPa as the elastic tissue modulus, a Poisson’s ratio of 0.3 and a linear hardening modulus of 0.05E [27] for the finite deformation-based plasticity model [28], [29]. This model is assigned to tolerate net section yielding with plasticity analysis assuming stress concentration in which the local stresses exceed the yield limit of a material with a nominally elastic region. Thus, in order to make accurate predictions related to the trabecular behaviour in vivo, this plasticity based model is required. The validity and comparison of the model was made with Fatihhi et al. [3] and all the constitutive parameters for fatigue were taken from this previous study.

The limitations presented by experimental analyses can be overcome by the use of computational method. Furthermore, the simulation is non-destructive and allows for better representation of localized trabecular level as variation in the properties of model can be reduced. In this study, FE analysis to investigate the behaviour of bovine trabecular bone model under fatigue simulation was first conducted to validate the assigned parameters and its subsequent fatigue response with the recorded data from experimental analysis [3]. The investigation was extended to evaluate the effect of off-axis loading on the fatigue behaviour of the trabecular bone model. Then the dependency of fatigue behaviour on the morphology of model was quantified.

Response of trabecular bone mechanical anisotropy towards different orientation has been quantified previously [27], [30], together with a few other authors who described the morphology and mechanical behaviour relationship of trabecular bone materials [5], [31]–[33]. However, huge variation in experimental data limits the accuracy of the results. This variation can be reduced significantly by restriction in number of samples, which subsequently reduces inter-specimen variation. This can be done by computational simulation and the model was developed with three different orientation as shown in Figure 2.

**Figure 2:**
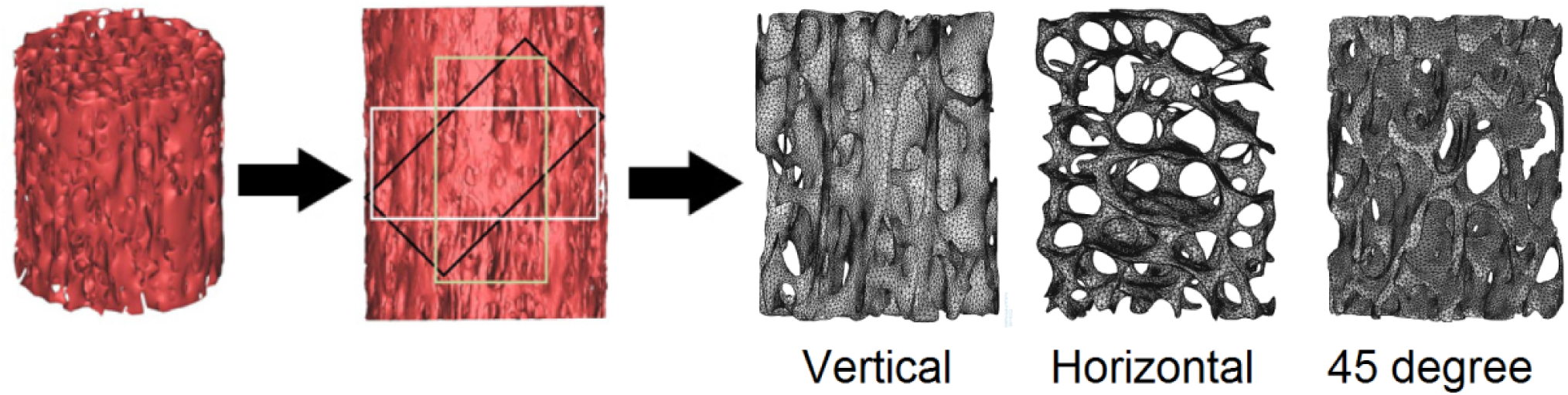
Three different models extracted from whole sample with different orientations: vertical (yellow), horizontal (white) and 45 degree (black).

## 3. Results and Discussion

Figure 3 shows the typical linear relationship of bone volume fraction and SMI with cycles to failure in both uniaxial and multiaxial loading. Good correlation in both BV/TV (*R*^2^ = 0.78 under uniaxial loading, and *R*^2^ = 0.68 under multiaxial loading) and SMI (*R*^2^ = 0.90 under uniaxial loading, and *R*^2^ = 0.86 under multiaxial loading) with cycles to failure was found. Satisfactory density of the trabecular bone structure ensure sufficient loading exertion. Furthermore, propagation of damage of the trabecular bone structure before failure is restricted by its plate-like structure, thus avoiding premature failure. However, no correlation between anisotropy in both type of loading for all trabecular orientation. Thus, the properties of the trabecular bone under multiaxial loading may be dependent on the individual struts, rather than the whole structure [2], [3]. Increase of cycles to failure is associated with high volume fraction and low SMI.

**Figure 3:**
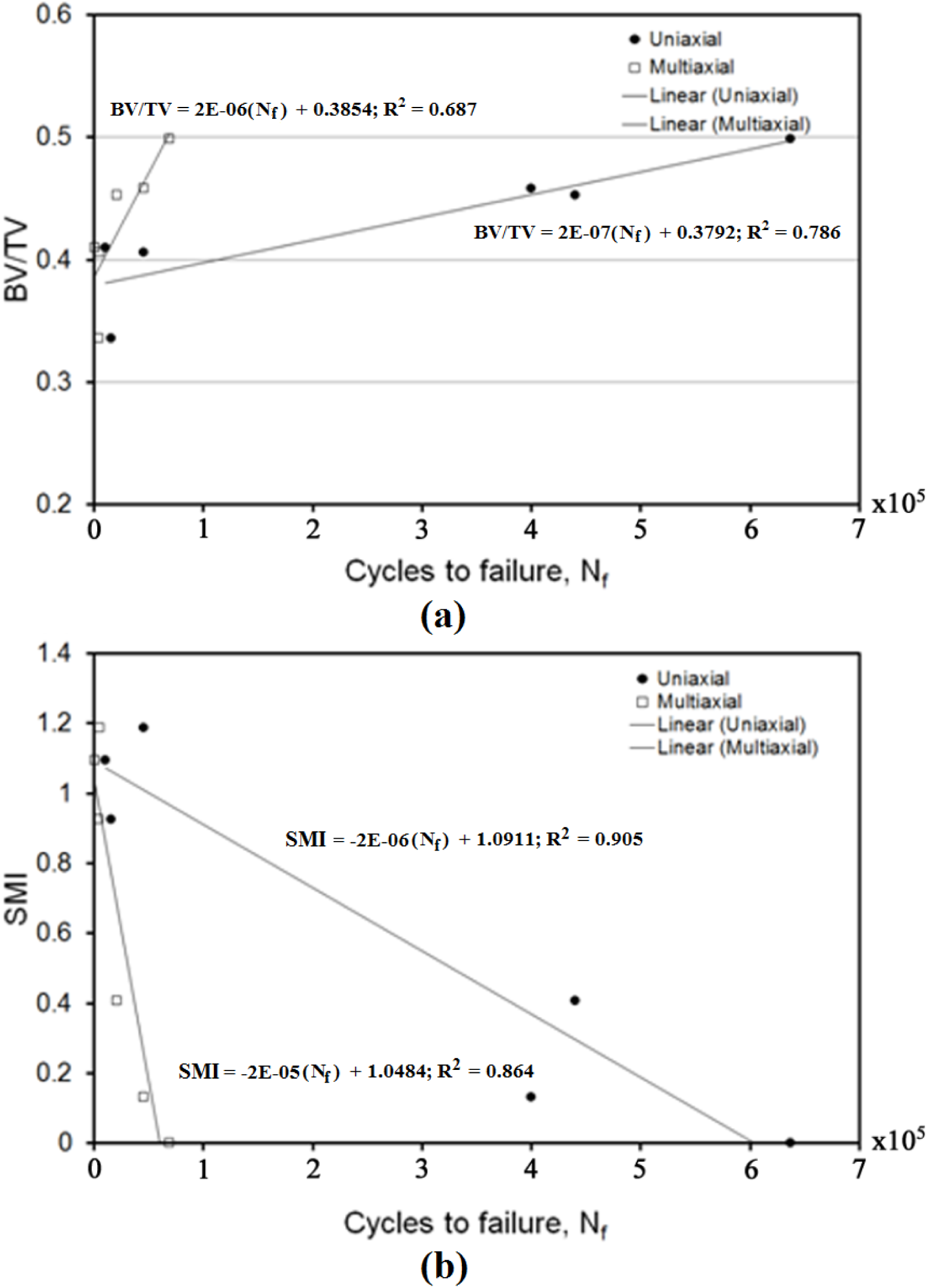
Linear relationship between (a) BV/TV and (b) SMI with cycles to failure, *N*_f_.

Regression analysis for other microarchitectural parameters is tabulated in Table 1 and Table 2 with coefficients of determination and *p*-values for cycles to failure and effective plastic strain in order to determine the contribution factor towards trabecular architecture or orientation with fatigue properties. All the coefficient values for microarchitectural parameters and their relative differences in cycles to failure and effective plastic strain were recorded. In most microarchitectural parameters such as BV/TV, SMI, Tb.Th, and Conn.D, samples under uniaxial loading showed better correlation (*R*^2^ > 0.50) with the cycles to failure, compared to samples under multiaxial loading. This is due to the stress distribution during loading. Uniaxial loading exerts a more even stress distribution compared to multiaxial loading, in which the distribution of stress are random and scattered. In contrast, Tb.Sp and DA of the samples under multiaxial loading demonstrated good correlation with the cycles to failure (*R*^2^ > 0.50). The trabecular bone samples are considered anisotropic, thus responded differently with different loading direction. An inverse correlation is observed in the relation of Tb.Th and SMI with the effective plastic strain. Thick trabecular struts resisted axial loading better than thin trabecular struts.

**Table 1:** Microstructural parameters of trabecular bone samples with coefficient of determination (*R*^2^) and *p*-value in relation to cycles to failure, *N*_f_ under both uniaxial and multiaxial loading conditions. Significant level with *p*-value < 0.05.

| Parameter | Uniaxial |  | Multiaxial |  |
| --- | --- | --- | --- | --- |
| | $R^2$ | $p$-value | $R^2$ | $p$-value |
| BV/TV | 0.786 | 0.064 | 0.687 | 0.131 |
| Tb.Th (mm) | 0.602 | 0.206 | 0.368 | 0.473 |
| Tb.Sp (mm) | 0.694 | 0.126 | 0.658 | 0.155 |
| BS/BV | 0.399 | 0.433 | 0.116 | 0.827 |
| DA | 0.139 | 0.793 | 0.484 | 0.331 |
| Conn.D (1/mm<sup>3</sup>) | 0.590 | 0.217 | 0.344 | 0.505 |
| SMI | 0.905 | $<0.05$ | 0.771 | 0.073 |

**Table 2:** Coefficient of determination for microstructural parameters of trabecular bone samples and *p*-value relative to effective plastic strain in both uniaxial and multiaxial loading condition.

| Parameter | Uniaxial |  | Multiaxial |  |
| --- | --- | --- | --- | --- |
| | $R^2$ | $p$-value | $R^2$ | $p$-value |
| **BV/TV** | 0.414 | 0.414 | 0.324 | 0.531 |
| **Tb.Th (mm)** | 0.284 | 0.585 | 0.685 | 0.134 |
| **Tb.Sp (mm)** | 0.220 | 0.676 | 0.329 | 0.523 |
| **BS/BV** | 0.425 | 0.400 | 0.405 | 0.426 |
| **DA** | 0.111 | 0.834 | 0.132 | 0.803 |
| **Conn.D (1/mm<sup>3</sup>)** | 0.100 | 0.850 | 0.037 | 0.944 |
| **SMI** | 0.533 | 0.276 | 0.476 | 0.340 |

Simulation in off-axis orientation predicted the fatigue life reduction and increased of plastic strain of the trabecular bone models. Highly localized stress contributes to early cycles to failure thus shorter fatigue life is observed in models subjected to multiaxial loading compared to models under uniaxial loading. Under both loading modes, the lowest cycles to failure is depicted by model in horizontal orientation (Figure 4). The trabecular bone model in this orientation has low volume fraction with minimum thickness of the trabecular structure, as such demonstrated by the trabecular bone with age-related bone loss. Consequently, the trabecular bone with this structure is more susceptible to fractures, especially on the vertebrae, femoral neck, and the distal radius. The model Stress-strain field of the trabecular bone model was determined by FE analyses, in which the prediction of mechanical parameters and fatigue life is investigated.

**Figure 4:**
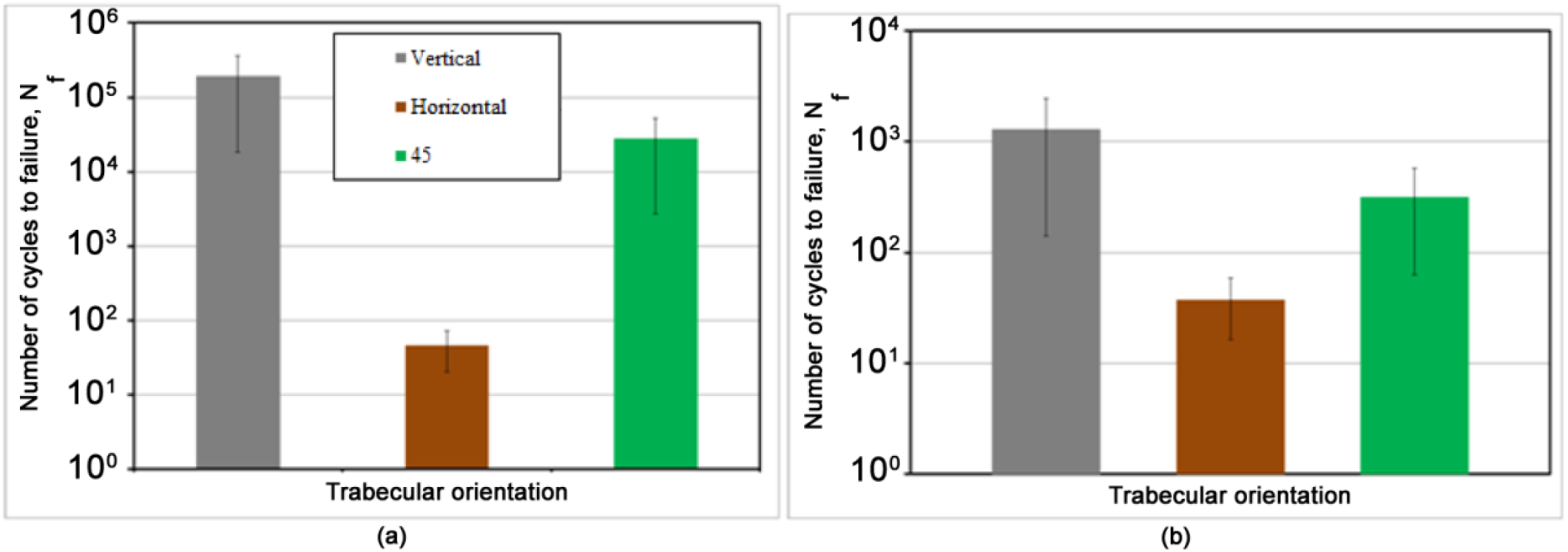
Comparison of (a) uniaxial and (b) multiaxial life prediction of the models at different trabecular orientation.

Furthermore, this study contributes to the evaluation of bone fracture risks under physiological loading conditions, in which the loading is not restricted to uniaxial loading. This is motivated by the fractures reported have been recognised to be initiated at the trabecular scale with high stress or strain [34], [35]. Besides, it is known that during habitual loading, the biological response is triggered by the mechanical stress [36], [37].

Von-mises stress distributions on the trabecular bone at three different orientations were plotted in Figure 5. Uniform stress distribution can be observed in vertical model with minor differences (ranging from 2.54 to 20.44 MPa) (Figure 5 (a)). More stress concentration is observed in model at 45° (ranging from 4.56 to 29.75 MPa) especially on the rod-like trabeculae due to non-uniformity of loading transition (Figure 5 (c)). On the other hand, horizontal model imitated severe osteoporotic trabecular structure with high stress amplitude (ranging from 28.43 MPa to 89.70 MPa) (Figure 5 (b)). This proves the weakness of osteoporotic bone relative to the off-axis loading.

**Figure 5:**
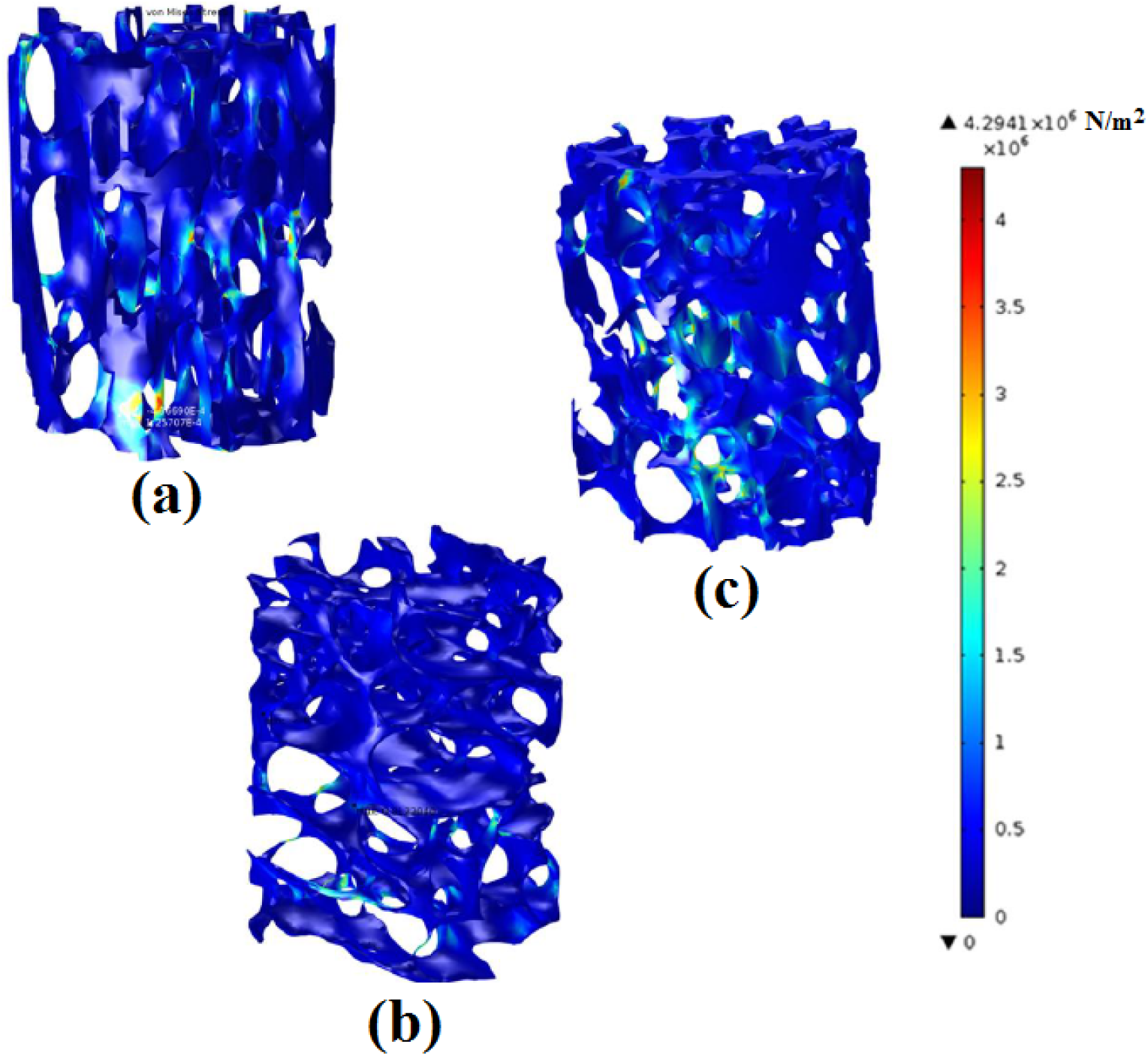
Difference in von-Mises stress distribution of trabecular surface corresponding to trabecular orientations; (a) vertical (b) horizontal, and (c) 45° subjected to uniaxial loading.

Under multiaxial loading, stress mapping of the trabecular model demonstrated different trend compared to uniaxial loading. Stress magnitude of the model in vertical orientation ranging from 5.85 to 24.90 MPa, with only small variation in stress distribution on trabecular as shown in Figure 6 (a). The 45° model demonstrates higher range of stress magnitude (8.95 to 44.66 MPa) than the vertical model, while model in horizontal orientation exhibited stress magnitude range of 8.02 to 55.98 MPa, exceeding both other models at different orientation. Main contribution to this distribution is due to the highest oblique orientation of trabeculae with strong interconnected several branches that resist bending upon the loading applied.

**Figure 6:**
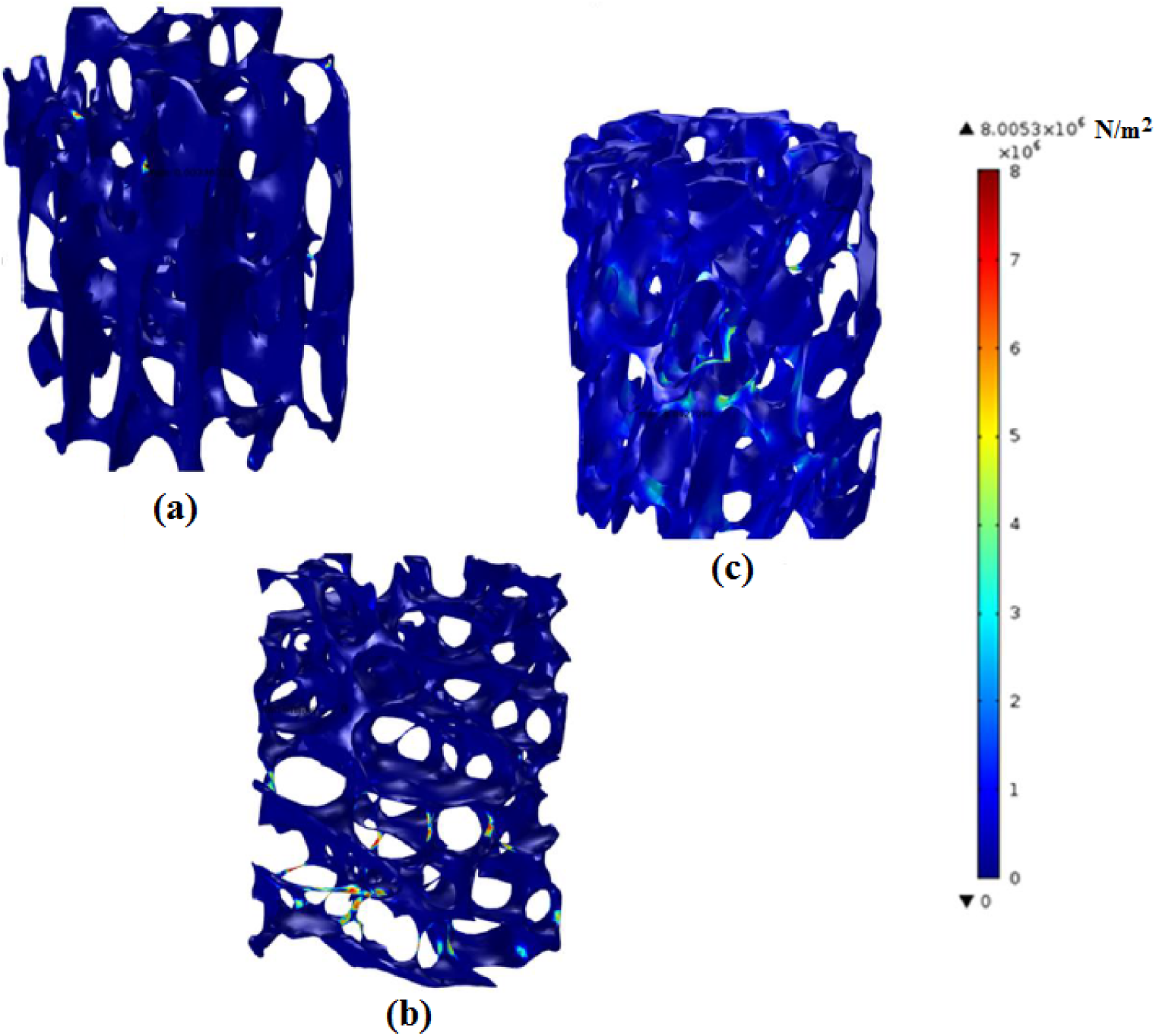
Difference in von-Mises stress distribution of the trabecular bone surface corresponding to trabecular bone orientations; (a) vertical (b) horizontal, and (c) 45° subjected to multiaxial loading.

The variation in the effective plastic strain across models at different orientation subjected to multiaxial loading is shown in Figure 7. The highest magnitude of strain is demonstrated by the model at horizontal orientation, while the model at 45° orientation shows magnitude different of about 1.6 times higher than that of the vertical model. Off-axis angle orientation reduces the capability of the trabecular structure to put up with stress during fatigue, thus reduction in fatigue life can be observed during experimental analysis [5]. In particular, present work simulated the same behaviour of trabecular structure by comparing three different trabecular models orientation. While increment in plastic strain of models in vertical and 45° orientation is found negligible, higher prominent strain is demonstrated by model in horizontal orientation. This is due to the structure of the model in which dominated by thin struts that are susceptible to bending and buckling. The results yielded in present work are in conjunction with previous studies which reported weaker trabecular in transverse orientation with higher shear stress concentration and shorter fatigue life [4]. Similar behaviour has also been observed in anisotropic reinforced composites, in which the strength reduction at different fibres orientation is demonstrated [38]. The same observation was also found previously in which demonstrated whole bone sustainability towards higher loading in the main axis, thus different orientation reduced the ability to withstand the load [39]. Differences in strains at failure have also been reported to be influenced by geometry [5]. Here, direct essentialities are true of providing detailed information on bone properties for improvement in fixation system.

**Figure 7:**
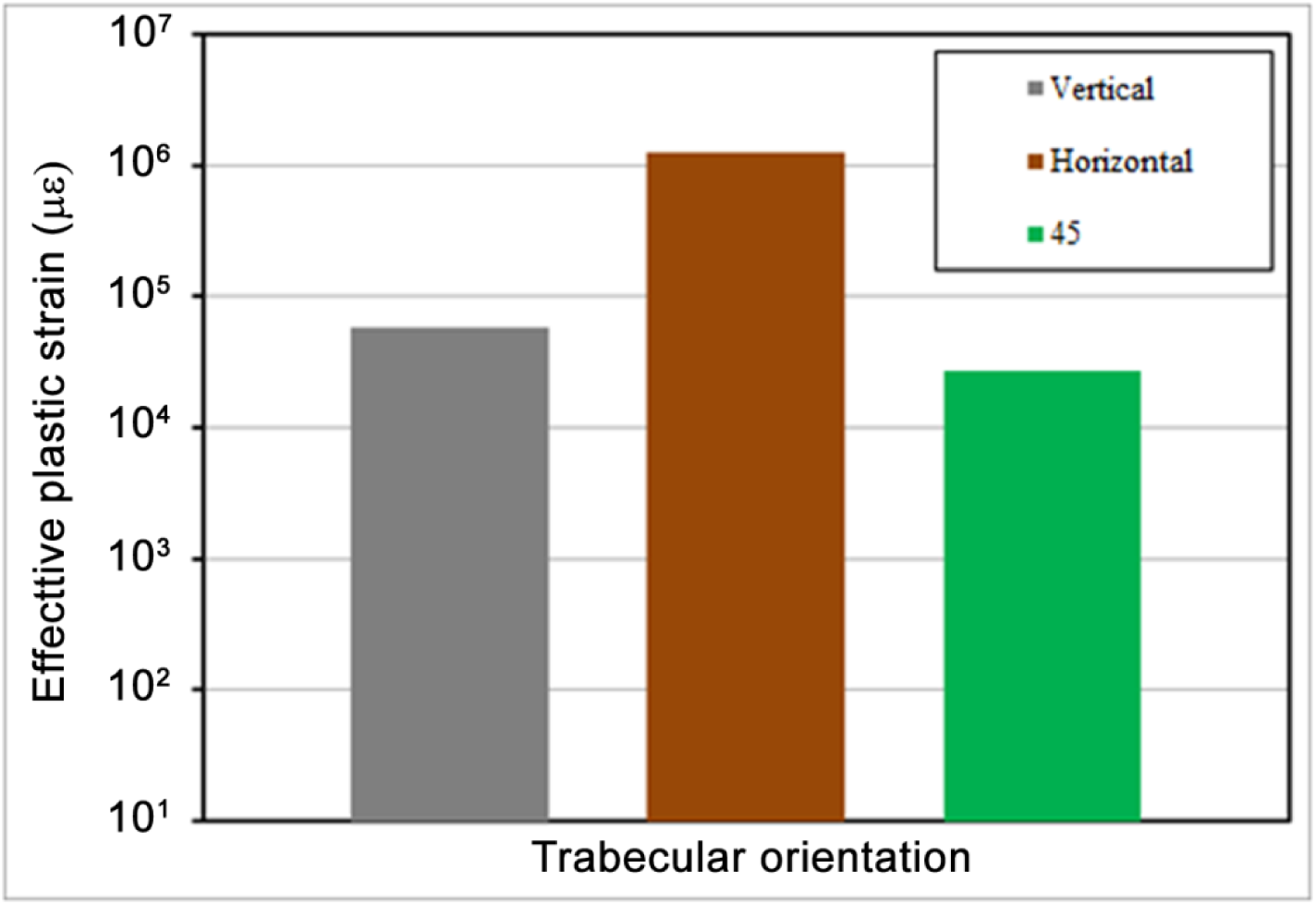
Effective plastic strain values for the corresponding trabecular models at different orientation.

## 4. Conclusion

Dramatic increased in effective plastic strain and decrease in cycles to failure was demonstrated by the trabecular bone models under multiaxial loading condition. Lower BV/TV, Tb.Th, and Conn.D resulted in lower number of cycles to failure, regardless of the loading conditions. However, the number of cycles to failure was found to be negatively correlated to the value of SMI. The reduction of fatigue life was five times lower in multiaxial condition in comparison to the fatigue life under uniaxial loading for all orientation. Additionally, off-axis orientation effect on the fatigue life of the trabecular bone was demonstrated the worst in horizontal trabecular bone model. Effective plastic strain was recorded the highest in horizontal model, while the model at 45° demonstrated 1.6 times higher effective plastic strain than the vertical ones.

